# Temperature-induced membrane trafficking drives antibody delivery to the brain

**DOI:** 10.1101/2024.10.04.616673

**Authors:** Fusheng Du, Qi Wan, Oleg O. Glebov

## Abstract

Despite quotidian occurrence of fever and hyperthermia, cell biological mechanisms underlying their effects remain unclear. Neurological complications of severe (>40°C) fever have been associated with increased blood-brain barrier (BBB) permeability due to structural disruption, while little is known about brain physiology of moderate fever. Here, we show that a temperature increase to 39-40°C increased fluid-phase uptake in PC12 cells and primary neurons. Uptake of selective cargoes showed that clathrin-mediated endocytosis and macropinocytosis were induced in a translation-dependent manner, consistent with a role for heat shock response. Exocytic recycling was also increased by hyperthermia, suggesting a comprehensive boost of membrane trafficking. Mild (<39°C) whole-body hyperthermia *in vivo* triggered fluid-phase uptake in various organs, notably enabling brain accumulation of an intravenously injected antibody that was blocked by dynamin inhibition. Taken together, our findings show that fever systemically regulates membrane trafficking, reveal dynamin-dependent endocytosis as a cell biological mechanism for temperature control of BBB permeability, and demonstrate a clinical potential of mild hyperthermia for facilitating brain delivery of biologic drugs.

**Highlights:**

- Temperature increase of 2-3°C upregulates endocytosis and exocytosis
- Temperature-induced upregulation of membrane trafficking requires protein translation and dynamin function
- Mild whole-body hyperthermia enhances fluid-phase endocytosis across the body
- Mild hyperthermia enables delivery of an exogenous antibody from the bloodstream into the brain through a dynamin-dependent pathway

## Introduction

Heat profoundly affects the functioning of the living matter through the commonly encountered phenomena of fever and hyperthermia. Fever constitutes an evolutionarily conserved response to injury and infection that is centred around controlled increase in body temperature, believed to confer protection against pathogens,^1–3^ while hyperthermia refers to uncontrolled increase in body temperature due to heat accumulation.^4,5^ Severe (>40°C) fever and hyperthermia are medical emergencies, presenting a rising burden on global public health due to the cumulative effects of antibiotic resistance and climate change.^2–4,6–14^ Most cases of fever and hyperthermia however feature mild or moderate temperatures up to 39-40°C, which may be associated with certain physiological and clinical benefits.^15–17^ Despite the prominence of fever and hyperthermia in and out of the clinic,^18,19^ the cell biological effectors of temperature on physiology remain poorly understood.

The brain is particularly affected by fever and hyperthermia, owing to its environmental sensitivity and metabolic demands. Temperatures above 40°C are associated with structural damage to the brain, in particular affecting the blood-brain barrier (BBB) - a specialised layer of endothelial cells separating the bloodstream from the brain.^3,7–9,9–11,20–22^ Considering that delivery of macromolecules into the brain is normally severely restricted by the BBB,^23^ thermal treatment has a potential for aiding drug brain delivery, especially for large biologic drugs such as immunoglobulins that cannot efficiently cross the BBB under normal conditions.^24–26^ Implementation of this approach would however necessitate combining BBB permeabilization with tissue damage mitigation,^20,21,27–29^ warranting deeper understanding of the cell biology underlying the physiological impact of temperature on the brain.

Membrane trafficking pathway termed transcytosis represents a key cell biology mechanism for macromolecule delivery across the BBB, whereby cargo undergoes endocytosis from the bloodstream space and exocytosis into the brain milieu.^24,30^ Transcytosis is normally tightly controlled, allowing only limited transport of specific cargoes; non-specific transcytosis of various cargoes across the BBB, for instance during aging,^31^ highlighting the potential for transcytosis to ferry macromolecules without overt BBB disruption. Based on early ultrastructural evidence^32^ and our recent findings demonstrating temperature effects on endocytosis in neurons,^33^ we hypothesised that temperature may regulate membrane trafficking in the brain; the work presented here provides evidence directly confirming this prediction.

## Results

To investigate the general link between temperature and membrane trafficking, we exposed rat pheochromocytoma 12 (PC12) cells to hypothermia (19-23°C) and hyperthermia (39-40°C) conditions. Calcein-AM/PI staining showed that neither hypothermia nor hyperthermia significantly affected cell viability (**Figure S1A, B**). Three hours of hyperthermia led to a robust increase in uptake of a fluid phase marker 4 kDa FITC-dextran, while hypothermia had no effect (**Figure 1A, B**). Similar effect as observed for a structurally unrelated fluorescent fluid-phase marker dye Lucifer yellow (**Figure S1C, D**). Immunostaining showed that hyperthermia induced a decrease in staining for an early endosomal marker EEA1 and increased staining for a late endosomal marker LAMP1, while hypothermia only decreased EEA1 staining (**Figure 1C-E**), showing that the increase in endocytosis was not coupled to the capacity of endosomal compartments. Hyperthermia but not hypothermia also increased F-actin levels (**Figure 1F, G**), consistent with the role for actin dynamics in endocytosis.^34^ Based on these observations, we sought to investigate hyperthermia-induced membrane trafficking in more detail.

**Figure 1.**
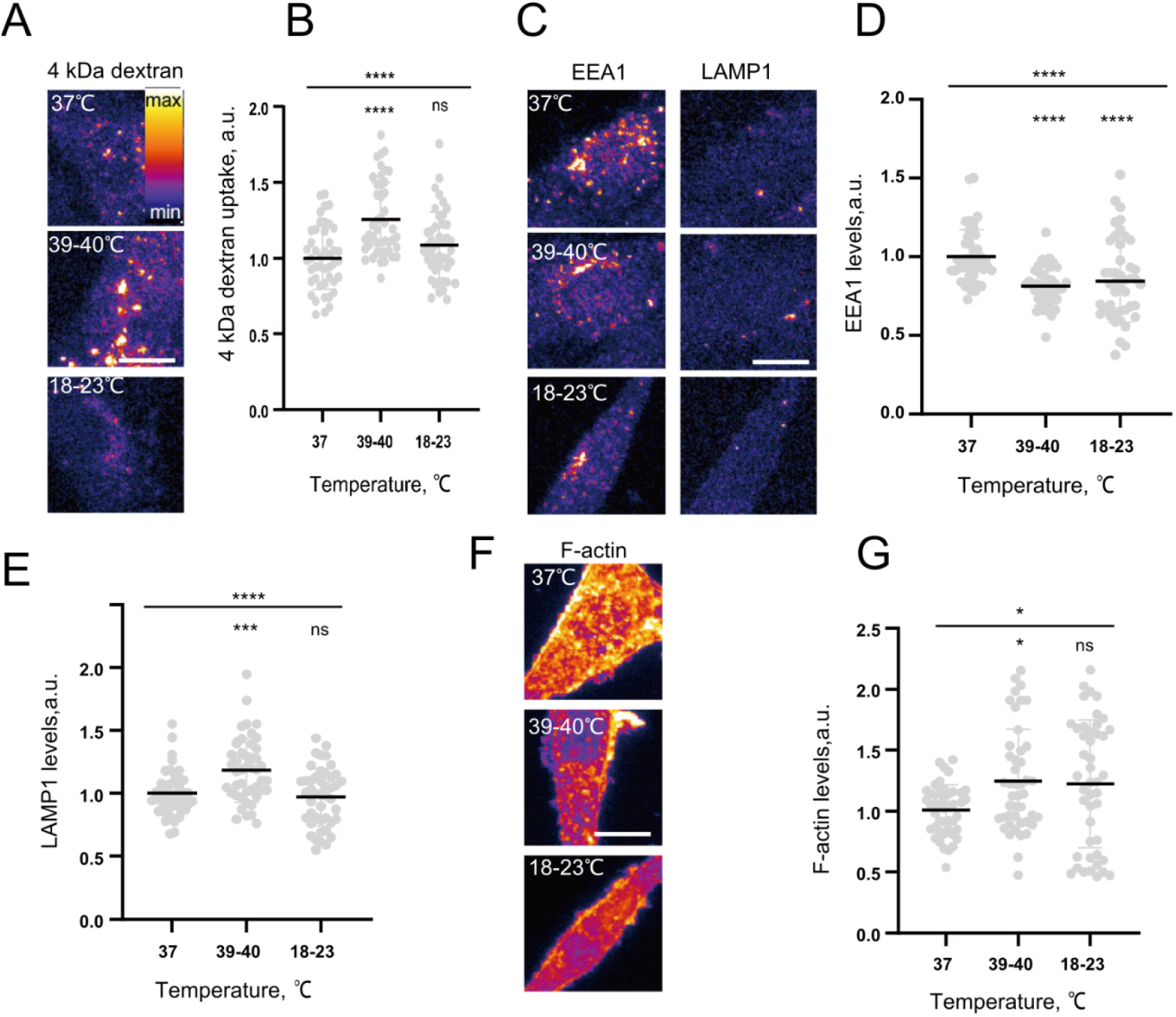
Hyperthermia induces fluid-phase uptake in cultured cells. (A) Representative images of 4kDa FITC-dextran internalisation in PC12 cells subjected to indicated temperatures for 3 hours. (B) Quantification of 4kDa FITC-dextran internalisation in PC12 cells treated with different temperatures for 3 hours. ****P < 0.0001, ns-not significant, Kruskal–Wallis test with Dunn’s multiple comparisons test. (C) Representative images of PC12 cells treated with different temperatures for 3 hours and immunostained for EEA1 and LAMP1. (D) Quantification of EEA1 levels in PC12 cells treated with different temperatures for 3 hours. ****P < 0.0001, Kruskal–Wallis test with Dunn’s multiple comparisons test. (F) Quantification of LAMP1 levels in PC12 cells treated with different temperatures for 3 hours. ****P < 0.0001,***P < 0.001, ns-not significant, Kruskal–Wallis test with Dunn’s multiple comparisons test. (G) Representative images of PC12 cells treated with different temperatures for 3 hours and stained for F-actin. (G) Quantification of F-actin levels in PC12 cells treated with different temperatures for 3 hours. *P < 0.05, ns-not significant, Kruskal–Wallis test with Dunn’s multiple comparisons test.

To investigate the role of specific pathways involved in hyperthermia-induced endocytosis, we focused on the two key endocytic pathways in PC12 cells, namely clathrin-mediated endocytosis (CME) involved in uptake of many receptors and ligands, and macropinocytosis providing internalisation of large fluid volumes and high molecular weight cargoes.^35^ We found that hyperthermia increased uptake of a FITC-labelled canonical CME cargo protein transferrin (FITC-Tf),^36^ and this uptake was inhibited by pharmacological blockade of a key CME regulatory protein dynamin by an antidepressant drug sertraline^37^ (**Figure 2A,B**), confirming the upregulation of CME. Uptake of a selective macropinocytosis marker 70 kDa TRITC-dextran^38^ was also enhanced following hyperthermia, while application of a macropinocytosis blocker 5-(n-ethyl-n-isopropyl)-amiloride (EIPA) prevented this increase (**Figure 2C, D**). Hyperthermia-induced increase in signal was observed for an even larger probe 2MDa FITC-dextran (**Figure S2A, B**). Uptake of 70 kDa TRITC-dextran but not FITC-Tf was rapidly induced by temperature, suggesting different dynamics (**Figure S1E-G**). Taken together, these results indicate that hyperthermia upregulates both CME and macropinocytosis in PC12 cells.

**Figure 2.**
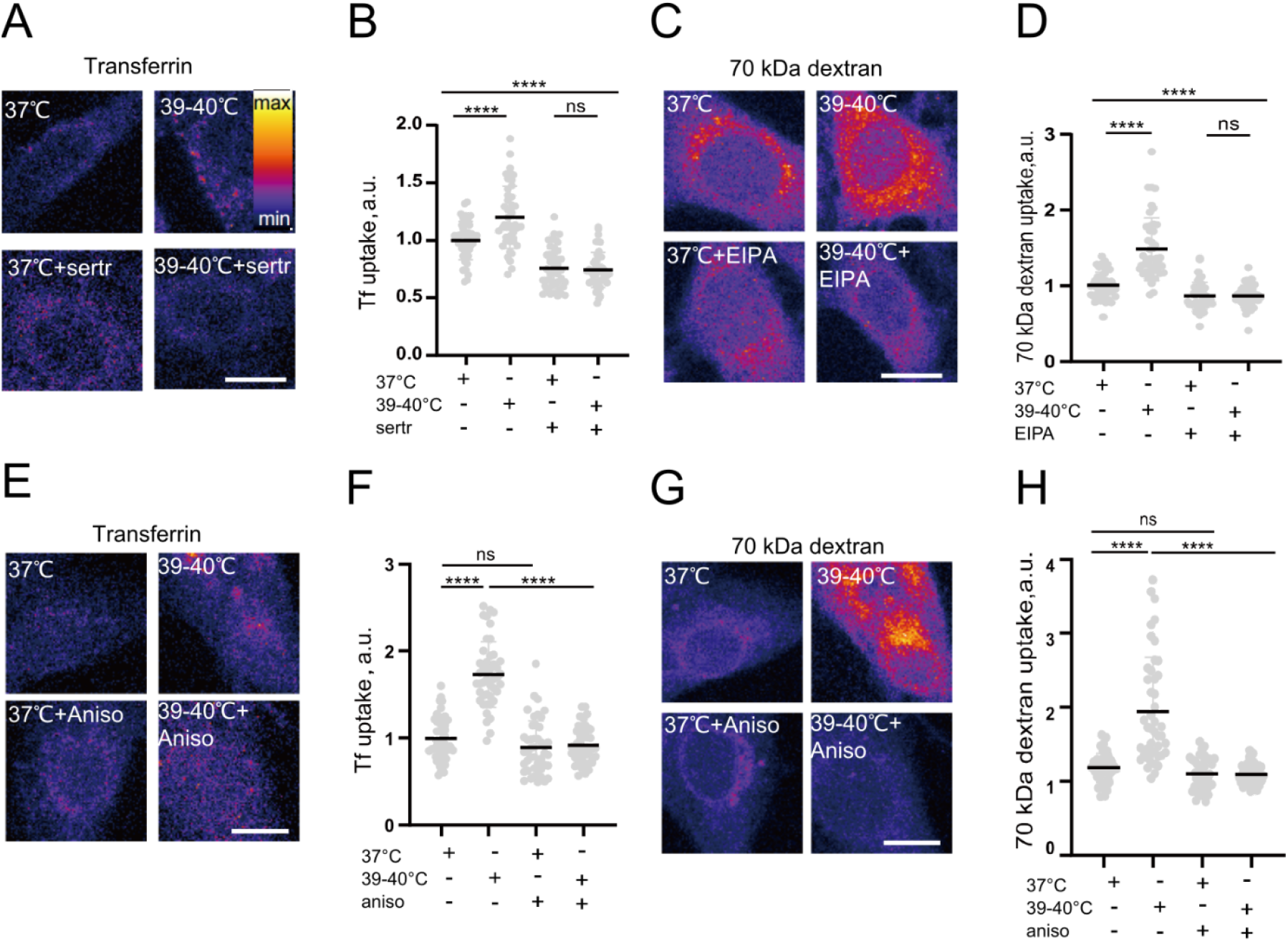
Hyperthermia induces CME and macropinocytosis in a translation-dependent manner. (A) Representative images of FITC-Tf internalisation in PC12 cells with 20μM sertraline for 15 min following 3 hours of hyperthermia. (B) Quantification of FITC-Tf internalisation in PC12 cells with 20 μM sertraline for 15 min after hyperthermia for 3 hours. ****P < 0.0001, ns-not significant, Kruskal–Wallis test with Dunn’s multiple comparisons test. (C) Representative images of 70 kDa TRITC-dextran internalisation in PC12 cells with 25μM EIPA after hyperthermia for 3 hours. (D) Quantification of PC12 cells incubated with 25μM EIPA after hyperthermia for 3 hours. ****P < 0.0001, ns-not significant, Kruskal–Wallis test with Dunn’s multiple comparisons test. (E) Representative images of FITC-Tf internalisation in PC12 cells treated with hyperthermia for 3 hours and anisomycin. (F) Quantification of FITC-Tf internalisation in PC12 cells treated with hyperthermia for 3 hours and anisomycin. ****P < 0.0001, ns-not significant, Kruskal–Wallis test with Dunn’s multiple comparisons test. (G) Representative images of 70kDa TRITC-dextran internalisation in PC12 cells treated with hyperthermia for 3 hours and anisomycin. (H) Quantification of 70kDa TRITC-dextran internalisation in PC12 cells treated with hyperthermia for 3 hours and anisomycin. ****P < 0.0001, ns-not significant, Kruskal–Wallis test with Dunn’s multiple comparisons test.

For further mechanistic understanding of hyperthermia-induced endocytosis, we evaluated the role for heat shock response, an evolutionarily conserved mechanism involving translation of heat shock proteins ^39,40^. Translation blocker anisomycin abolished hyperthermia-induced uptake of FITC-Tf (**Figure 2E, F**) and 70 kDa TRITC-dextran (**Figure 2G, H**). Immunostaining showed an increase in levels of the canonical heat shock proteins Hsp70 and Hsc70 that was also abolished by anisomycin treatment (**Figure S2C-F)**, confirming that 39-40°C hyperthermia was sufficient to induce the heat shock response and reaffirming the role of heat-shock response in hyperthermia-induced membrane trafficking. To investigate the effects of hyperthermia on membrane trafficking in differentiated polarised cells, we quantified uptake in primary neurons. Much like in PC12 cells, hyperthermia significantly increased endocytosis of both FITC-Tf and 70 kDa TRITC-dextran in primary neurons in a translation-dependent manner (**Figure S3A-C**). These data show that hyperthermia treatment increased endocytosis in polarised cell types.

To test the effects of hyperthermia on exocytosis, we measured loss of internalised signal from pre-loaded cargo signal over time.^36^ After 20 minutes, the rate of FITC-Tf signal loss in cells subjected to hyperthermia was significantly increased in comparison to the control cells, indicating that exocytic recycling of endocytosed FITC-Tf was more enhanced by hyperthermia than endocytosis (**Figure 3A, B**). For 70 kDa TRITC-dextran, signal loss rate in hyperthermia-treated cells was not significantly different from the control cells, indicating that hypothermia-induced increase in fluid-phase endocytosis was matched to that in endocytosis (**Figure 3C, D**). Taken together, these data show that hyperthermia enhances exocytosis as well as endocytosis.

**Figure 3.**
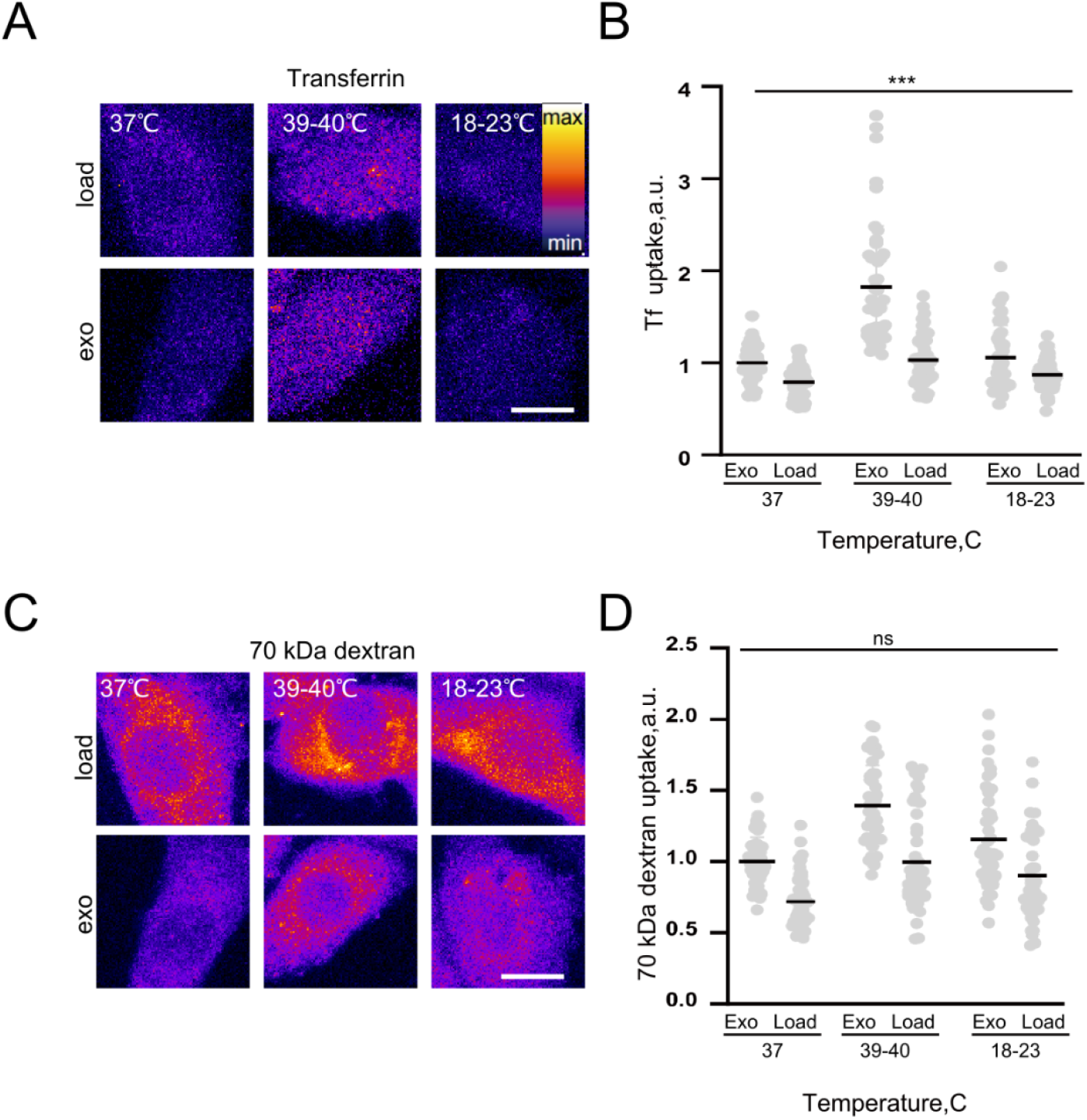
Hyperthermia induces exocytosis. (A) Representative images of FITC-Tf exocytosis in PC12 cells treated with different temperatures. Load - cells loaded with Tf for 1 hour, exo - cells loaded with Tf for 1 hour following 20 min in Tf-free medium. (B) Quantification of FITC-Tf exocytosis in PC12 cells treated with different temperatures. ***P < 0.001 for the effect of temperature, 2-way ANOVA. (C) Representative images of 70 kDa TRITC-dextran exocytosis in PC12 cells treated with different temperatures. Load - cells loaded with Tf for 1 hour, exo - cells loaded with Tf for 1 hour following 20 min in Tf-free medium. (D) Quantification of 70 kDa TRITC-dextran exocytosis in PC12 cells treated with different temperatures. ns, no significant effect of temperature, 2-way ANOVA. N ≥ 3 independent experiments, n ≥ 45 cells. Scale bar, 10 μm.

On the basis of the above evidence for hyperthermia-induced membrane trafficking in cultured cells, we sought to establish its role in transport across biological barriers *in vivo*. To this end, we induced whole-body hyperthermia (WBH) in mice^41^ and measured penetration of distally administered markers 4 kDa FITC-dextran and Evans blue into various organs as described by us recently.^36^ The WBH protocol induced a temperature increase to 38.3-39°C within 3 hours (**Figure 4A, B**), equivalent to mild fever and well below the temperature range associated with structural damage to the tissue. We found a significant increase in 4 kDa FITC-dextran accumulation for lung and heart, and a trend towards increased accumulation in other organs including the brain (**Figure 4C, 4D, S4A, S4B**). While normothermic control animals showed no signs of brain permeation by Evans blue, WBH treatment induced significant Evans blue accumulation (**Figure 4C, E**), consistent with induction of BBB permeability by WBH.

**Figure 4.**
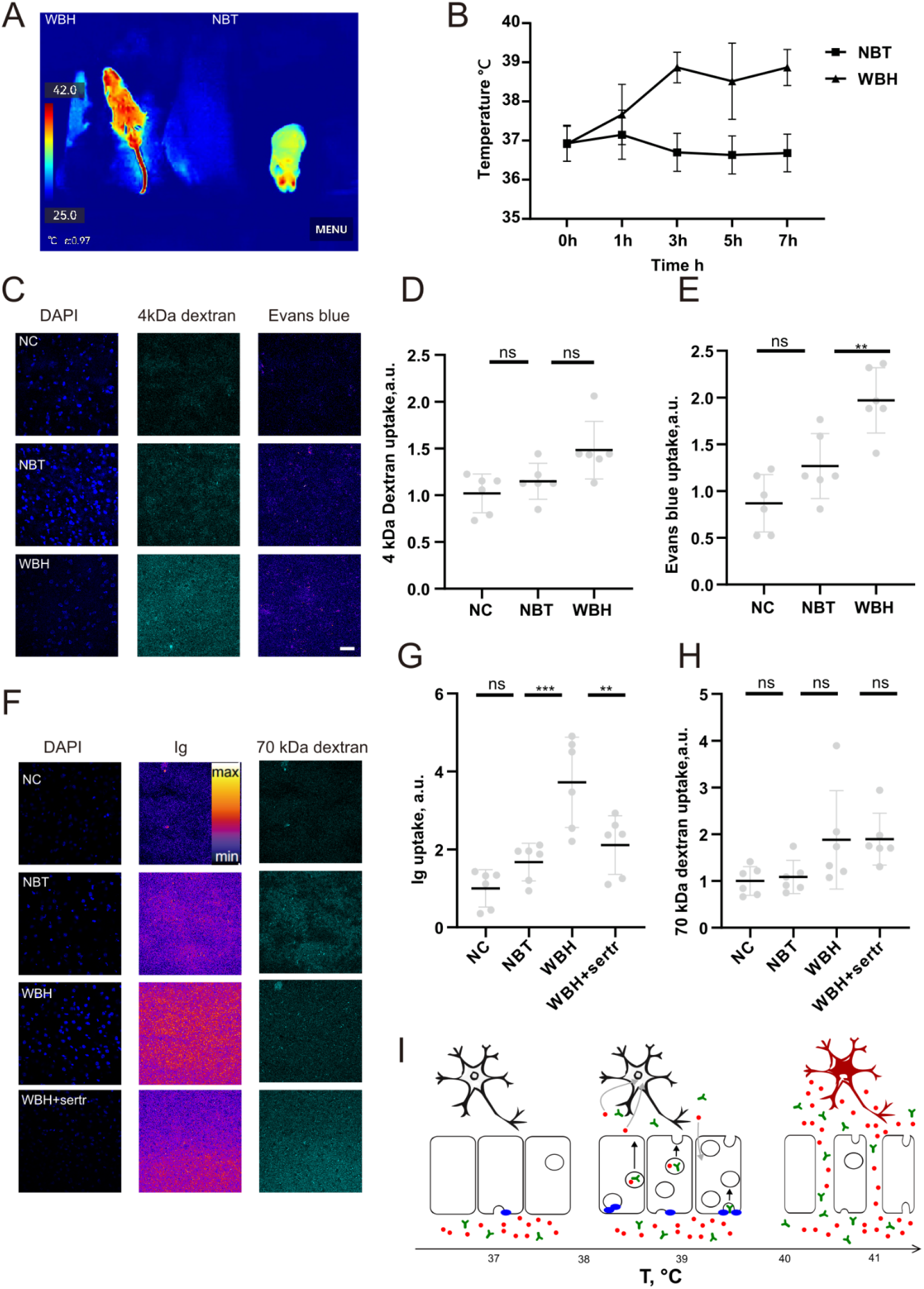
Transcytosis of exogenous antibody across the BBB into the brain induced by mild whole-body hyperthermia. (A) Representative illustration of hyperthermia induction in mice. Left, treated mouse; right, control mouse. WBH, whole body hyperthermia; NBT, normal body temperature. (B) Time course of WBH induction in mice. (C) Representative image of cortical brain sections from mice with injected 4 kDa FITC-dextran and Evans blue. NC, negative control (no injection). (D) Quantification of brain uptake of 4 kDa FITC-dextran. ns, not significant, 1-way ANOVA with Dunn’s multiple comparisons test. Data was normalised to the signal in NC. (E) Quantification of brain uptake of Evans blue. **P<0.01, 1-way ANOVA with Dunn’s multiple comparisons test. Data was normalised to the signal in NC. (F) Representative images of cortical brain sections from mice with injected Ig and 70 kDa dextran . (G) Quantification of brain uptake of exogenous immunoglobulin (Ig). ***P<0.001, **P<0.01, ns - not significant, 1-way ANOVA with Dunn’s multiple comparisons test. Data was normalised to the signal in NC. (H) Quantification of brain uptake of 70 Da TRITC-dextran. ns - not significant, 1-way ANOVA with Dunn’s multiple comparisons test. Data was normalised to the signal in NC. (I) A speculative model for physiological effects of fever/hyperthermia on permeability of the BBB. Ig (immunoglobulin, green) represents macromolecules of interest for brain delivery, glutamate (red) represents undesirable molecules to be excluded from the brain. At *normal body temperature (left)*, transcytosis through the BBB is restricted, preventing ingress of both Ig and glutamate into the brain space. Under conditions of *mild fever/hyperthermia (middle)*, membrane trafficking requiring activity of dynamin (blue) is induced, enabling transcellular route of Ig and possibly glutamate to the brain. Due to limited capacity of temperature-induced transcytosis, glutamate is effectively removed from the brain milieu through the action of glutamate transporters, curbing excitotoxicity. In *severe fever/hyperthermia (right)*, structural integrity of the BBB is compromised, resulting in uncontrolled entry of both Ig and glutamate into the brain, inducing excitotoxicity and cell death.

To further investigate the translational relevance of WBH-induced BBB permeability, we sought to test whether it can enable uptake of larger biologically relevant macromolecules and, if so, whether this process involves membrane trafficking. The two main membrane trafficking pathways for transcellular uptake are CME and caveolar endocytosis, both of which require dynamin function.^24^ We therefore investigated uptake of an immunoglobulin (Ig) and 70 kDa TRITC-dextran in presence of sertraline to block dynamin-dependent transcytosis. WBH robustly induced brain accumulation of Ig which was abolished by sertraline injection, indicating that Ig was was indeed transported in a dynamin-dependent manner (**Figure 4F, G**). Conversely, WBH did not result in significant uptake of 70 kDa TRITC-dextran, suggesting that macropinocytosis was not induced and/or that BBB integrity was unaffected (**Figure 4F, H**). Taken together, these results show that membrane trafficking induced by mild WBH requires dynamin function and can carry immunoglobulins from the bloodstream into the brain.

## Discussion

Our study provides a key mechanistic insight into the cell biology of fever and hyperthermia in the brain. To thi day, physiologically relevant temperature regimes of mild fever and hyperthermia in the brain have remained understudied, the focus of investigation being on decidedly pathological extremes.^4,5,8,11^ Our work addresses this gap in knowledge, showing that increases in temperature as little as 2°C can elicit salient changes in membrane trafficking, suggesting a simple model for effects of temperature on the BBB permeability (**Figure 4I**). More broadly, a common cell biological basis underlying the wide-ranging physiological impacts of febrile temperature range across the body is likely to have far-reaching implications, given the central role of membrane trafficking in health and disease.^30^

The physiological role of membrane trafficking in fever remains to be established. Induction of macropinocytosis reported here is consistent with the role of fever in enhancing phagocytosis and activation of lymphocytes^42,43^ and may upregulate other relevant pathways, e.g. at the immunological synapse^44,45^ or in fever-associated immune signalling pathways such as cytokine signalling.^46–48^ Further, membrane trafficking is likely to contribute to a wide range of fever-associated processes, including endocytosis of pathogens,^49^ processing of pathogens by relevant immune cells, cell migration, and extravasation. More generally, upregulation of membrane trafficking may provide a compensatory mechanism to counter the deleterious effects of temperature increase across the body, e.g. by stimulating lysosomal clearance of misfolded proteins.^50^ Conversely, an increase in temperature over a certain threshold may result in dysregulation of membrane trafficking, contributing to adverse effects of hyperthermia and high fever.^11^ Direct testing of these predictions will require manipulation of temperature-induced membrane trafficking, e.g. by dynamin inhibitors, in physiologically relevant experimental models of mild and severe fever.

Our data show that temperature-induced membrane trafficking is associated with expression of heat shock proteins and requires protein biosynthesis. This evidence is consistent with canonical heat shock response,^39^ rather than translation-independent processes such as cooling-induced synaptic enhancement^33^ or regulation of cytosolic viscosity^51^, yet a role for other temperature sensors *e*.*g*. ion channels cannot be explicitly ruled out ^52,53^. Mechanistically, heat shock proteins could regulate endocytosis via direct interaction between heat shock proteins and CME machinery ^54 55^, actin dynamics ^56^, and directly with biomembranes ^57,58^, or indirectly through protein phase separation ^59–61^. For now, the molecular mechanism coupling heat shock response with membrane trafficking remains unclear, warranting further investigation.

The key translationally relevant conclusion stemming from our work is that low-level hyperthermia induces wide-ranging fluid-phase endocytosis *in vivo*, sufficient to ferry therapeutically relevant macromolecules from the bloodstream into the brain. It is therefore conceivable that membrane trafficking under a low-febrile temperature regime may be translated into a generalisable drug delivery protocol for a wide variety of cargoes. Considering that low-to-moderate fever and hyperthermia is easy to control and unlikely to pose health risks,^1,11^ this alternative approach would obviate many safety and cost-efficiency concerns associated with delivery of biologics.^25,25,26,62–65^ Crucially, our experimental evidence directly confirms the validity of this approach for brain delivery of antibodies, the currently dominant class of biologics.^66^ Although it remains to be seen to what extent fever and hyperthermia increase BBB permeability in humans, in due course our findings may inform development of forward clinical trials towards temperature-aided drug delivery into the brain, as well as other hard-to-reach tissues of the body.

## Acknowledgements

The authors thank Dr E. Barker for comments on the draft manuscript. This work was funded by the Lewy Body Society (OOG2019/2020) and the National Natural Science Foundation of China (32070772).

## Methods

### Reagents, drugs, fluorescent markers

Cell culture materials were from Solarbio and Procell. Institute for Cancer Research (ICR) mice and Sprague-Dawley rats were from Jinan Pengyue Laboratory Animal Co. 4 kDa FITC-dextran and 2 MDa FITC-Dextran were from Sigma-Aldrich. 70 kDa TRITC-dextran was from Chondrex Inc. FITC-Transferrin was from Jackson Immunoresearch. Lucifer yellow was from Santa Cruz. Anisomycin and doxorubicin was from Macklin. EIPA was from Selleck. Evans blue was from Solarbio. Sertraline and Apoptozole was from AbMole. 647 phalloidin was from Yeasen Biotechnology. Calcein-AM/PI Double Staining Kit was from Meilun. Rabbit immunoglobulin (Ig) was from Synaptic Systems.

### Cell culture

PC12 cells were grown and maintained in high glucose Dulbecco’s Modified Eagle medium (DMEM) (Solarbio) containing 10% foetal bovine serum (FBS) (Procell/Gibco) and 1% penicillin/streptomycin (Gibco) at 37°C in a humidified incubator with 5% CO2. PC12 cells were seeded into 24-well culture plates (3×104 cells/well, Corning Costar) at a density of 50-60% for temperature treatment and subsequent cell uptake, immunocytochemistry and Phalloidin staining. Primary rat neuronal cultures were isolated and maintained according to the standard Banker method ^67^.

### Temperature treatment and membrane trafficking assays

After the PC12 cells had grown in 24-well plates and reached 50-60% confluence, they were used for experimentation. Cells were exposed to hypothermia (18-22°C), normothermia (37°C) and hyperthermia (39-40°C) for 3 hours, using dedicated cell incubators. Culture medium was high glucose DMEM (Solarbio) containing 20 mM HEPES, used at 500 μl per well. Same treatment was applied to neurons, except the culture medium was Neurobasal (Gibco) with B27 and glutamine.

After treatment at different temperatures for 3 hours, medium was removed and fresh medium containing fluorescently labelled compounds was added, whereupon cells were transferred to the 37°C incubator for uptake. After uptake, cells were fixed with 4% paraformaldehyde (PFA) and washed six times with phosphate buffer (PBS) for five minutes each. Slides were then mounted in Fluoromount-G mounting medium (Southern) and stored at 4°C until imaging. Final concentrations of 4 kDa FITC-dextran, 70 kDa TRITC-dextran, 2 MDa FITC-dextran, FITC-Tf, Lucifer yellow were 1 mg/mL, 1 mg/mL, 2 mg/mL, 5 μg/mL, 1 mg/mL respectively. EIPA was used to block macropinocytosis at a concentration of 25 μM, sertraline was used to block CME at a concentration of 20 μM, and anisomycin was used at 5 μg/mL.

To measure exocytosis, cells were washed three times with 500μl of medium and then loaded with medium containing 70 kDa FITC-dextran or FITC-Tf for 1 hour at indicated temperatures. Subsequently, cells were removed from the incubator, washed three times with 500 ul culture medium, and then either fixed immediately in 4% PFA (no exocytosis control) or incubated with new culture medium for 20 min at indicated temperatures and then fixed as above. Cells were mounted in Fluoromount-G and stored at 4°C until imaging.

### Whole body hyperthermia in vivo

ICR mice (8-10 weeks old, male, weighing 28-32 g) were randomly allocated into WBH, NBT, and NC groups. The WBH group was put into a heating box, and the temperature was increased by 0.5°C every 15 min to 39°C and maintained at this level for 6 hours. Once per 2 hour period, all mice were placed at room temperature for a 15 min rest, while rectal temperature was measured and 0.1mL saline was injected intraperitoneally. Rectal temperature was measured at a final rest for thermal imaging. NBT and NC groups were maintained at room temperature (18-22°C), with the same procedures for injection and temperature measurement as described above. All animals were sacrificed one hour later. The brains were fixed with 4% PFA, cut into into 25 or 30 μm thick sections with a frozen slicer, mounted onto positively-charged microscope slides in media containing 4′,6-diamidino-2-phenylindole (Solarbio), and imaged using a confocal microscope.

### Blood-brain barrier measurements in vivo

For measurement of blood-brain barrier permeability on Evans blue and 4 kDa FITC-dextran, Evans blue and 4 kDa FITC-dextran were dissolved in PBS and injected into the tail vein. The dosage of 4kDa FITC-dextran was 200 μg/kg, while Evans blue was administered as a 2% solution at 4 mL/kg. Equal volumes of PBS were injected into the NC group animals.

For measurement of blood-brain barrier permeability on Ig and 70 kDa TRITC-dextran and the effect of sertraline on its permeability. ICR mice were randomly allocated into groups as above plus the whole body hyperthermia group with sertraline (WBH+sertr). Sertraline was dissolved in PBS and injected intraperitoneally at dosage of 10 mg/kg; other groups were injected with an equal volume of PBS. Three hours after the sertraline injection, Ig and 70 kDa TRITC-dextran were dissolved in PBS and injected into the tail vein in dosages of 20 μg/kg and 200 μg/kg respectively. Temperature treatment and other operations were as above.

### Immunocytochemistry and phalloidin staining

After temperature treatment, cells were transferred to room temperature and washed once in PBS. Coverslips were fixed with 4% PFA in PBS for 20 min and afterwards washed six times in PBS. For blocking, coverslips were incubated with 0.3% Triton-X 100 in PBS supplemented with 5% goat serum for 60 min. Following the primary antibody incubation for 2 hours, coverslips were washed six times in PBS, and incubated with Alexa Fluor 488 and Alexa Fluor 594 conjugated secondary antibodies for 1 hour. After 6 more washes in PBS, coverslips were mounted onto slides in Fluoromount-G mounting medium and stored at 4°C until imaging. The antibodies used in this study are listed in **Table S1**. For staining with phalloidin, coverslips were fixed and blocked as described above, then incubated with 1/2000 phalloidin for 2 hours, washed six times in PBS, then mounted and stored as above.

### Confocal microscopy imaging

Samples were imaged on Nikon Eclipse Ti2 laser confocal microscope. The imaging systems were controlled by NIS Elements 2.0 software. The imaging parameters for Nikon Eclipse Ti2 were as follows. The following image acquisition settings were used for serial confocal z-stack images: cells - 1.5 μm step, 512 x 512 pixels, 1x zoom, 100x magnification; tissue sections - 0.5 μm step (short-term assay) or 1.5 μm (long-term assay) 512 x 512 pixels, 1x zoom, 40x magnification. Pinhole size was kept at a suitable (1-2AU) Airy unit. Excitation laser wavelengths were 488 nm, 561 nm and 647 nm. Bandpass filters were set at 500−550 nm (FITC, Alexa Fluor 488) and 570−620 nm (TRITC, AlexaFluor 594) and 650−750nm (Alexa Fluor 647). Gain, exposure and offset settings were optimised within each experiment to ensure appropriate dynamic range, low background and optimal signal/noise ratio.

### Statistical analysis

Statistical analysis was carried out using the GraphPad Prism v5 (https://www.graphpad.com/). Data distributions were assessed for normality using d’Agostino and Pearson omnibus normality tests. For normally distributed datasets, two sample t test, 1-way ANOVA, with Dunn’s, Bonferroni’s post tests and 2-way ANOVA were used to assess statistical significance as appropriate; for non-normally distributed datasets, Mann-Whitney rank test, Kruskal-Wallis test and Dunn’s post test were used. The data sets are presented in the form of scatter plots, bar plots and line plots, with plot averages as lines.

## Supplemental Materials

**Figure S1.**
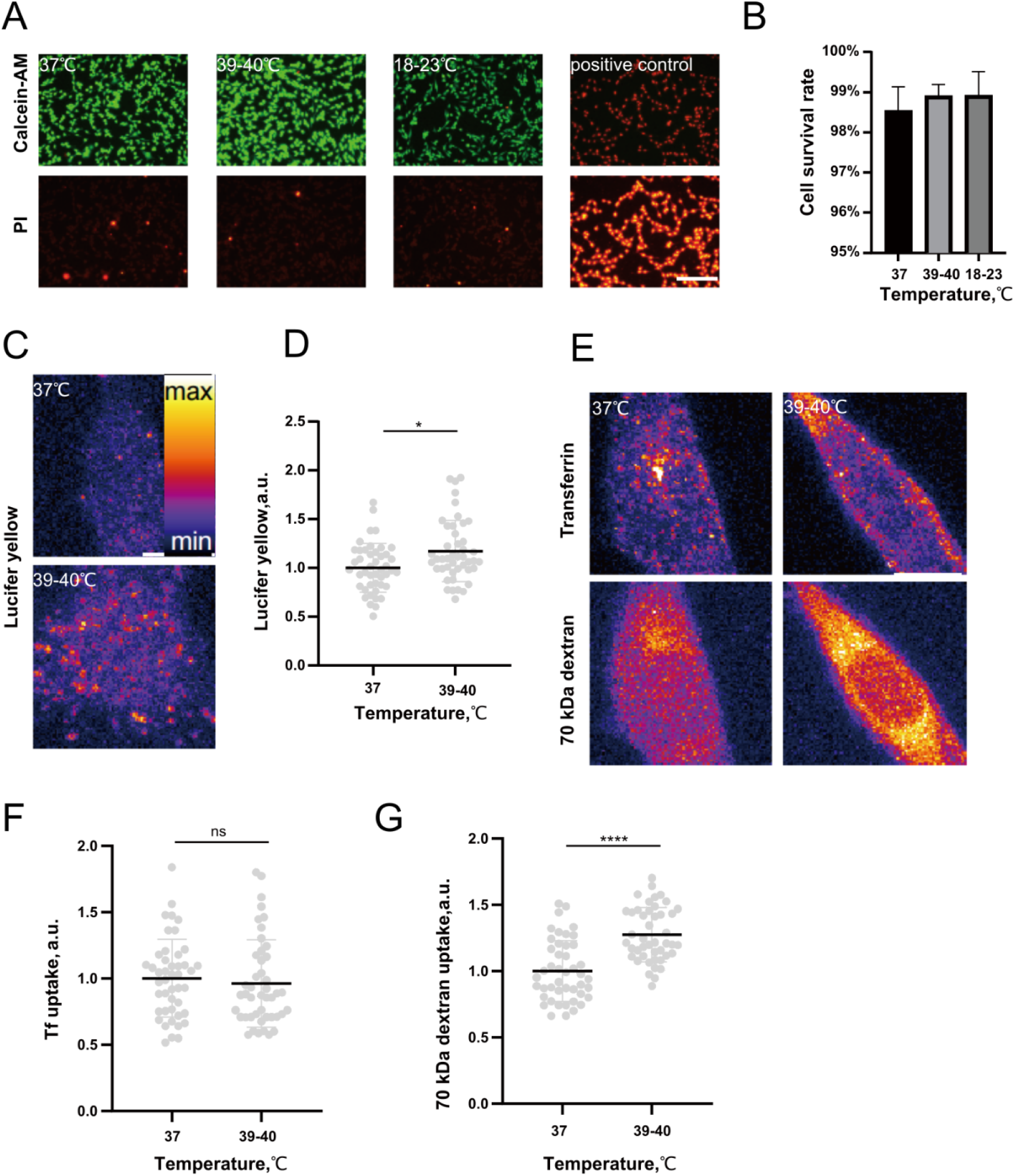
Further analysis of hyperthermia-induced endocytosis. (A) Representative images of Calcein-AM/PI double stained in PC12 cells subjected to different temperatures for 3 hours. Scale bar, 200 μm. (B) Rate of cell survival in each image of PC12 cells subjected to different temperatures for 3 hours. (C) Representative images of Lucifer yellow internalisation in PC12 cells subjected to hyperthermia for 3 hours. N ≥ 3 independent experiments, n ≥ 45 cells. Scale bar, 10 μm. (D) Quantification of Lucifer yellow endocytosis.*P < 0.05, Mann–Whitney test. (E) Representative images of PC12 cells uptake of FITC-Tf and 70 kD TFITC-dextran for 15 min at 37°C vs at 39-40°C without preincubation. N = 3 independent experiments, n = 45 cells. Scale bar, 10 μm. (F) Quantification of uptake of PC12 cells uptake of FITC-Tf for 15 min at 37°C vs 39-40°C without preincubation. ns-not significant, Mann–Whitney test. (G) Quantification of uptake of PC12 cells uptake 70 kDa TRITC-dextran for 15 min at 37°C vs at 39-40°C without preincubation. ****P < 0.0001, Mann–Whitney test.

**Figure S2.**
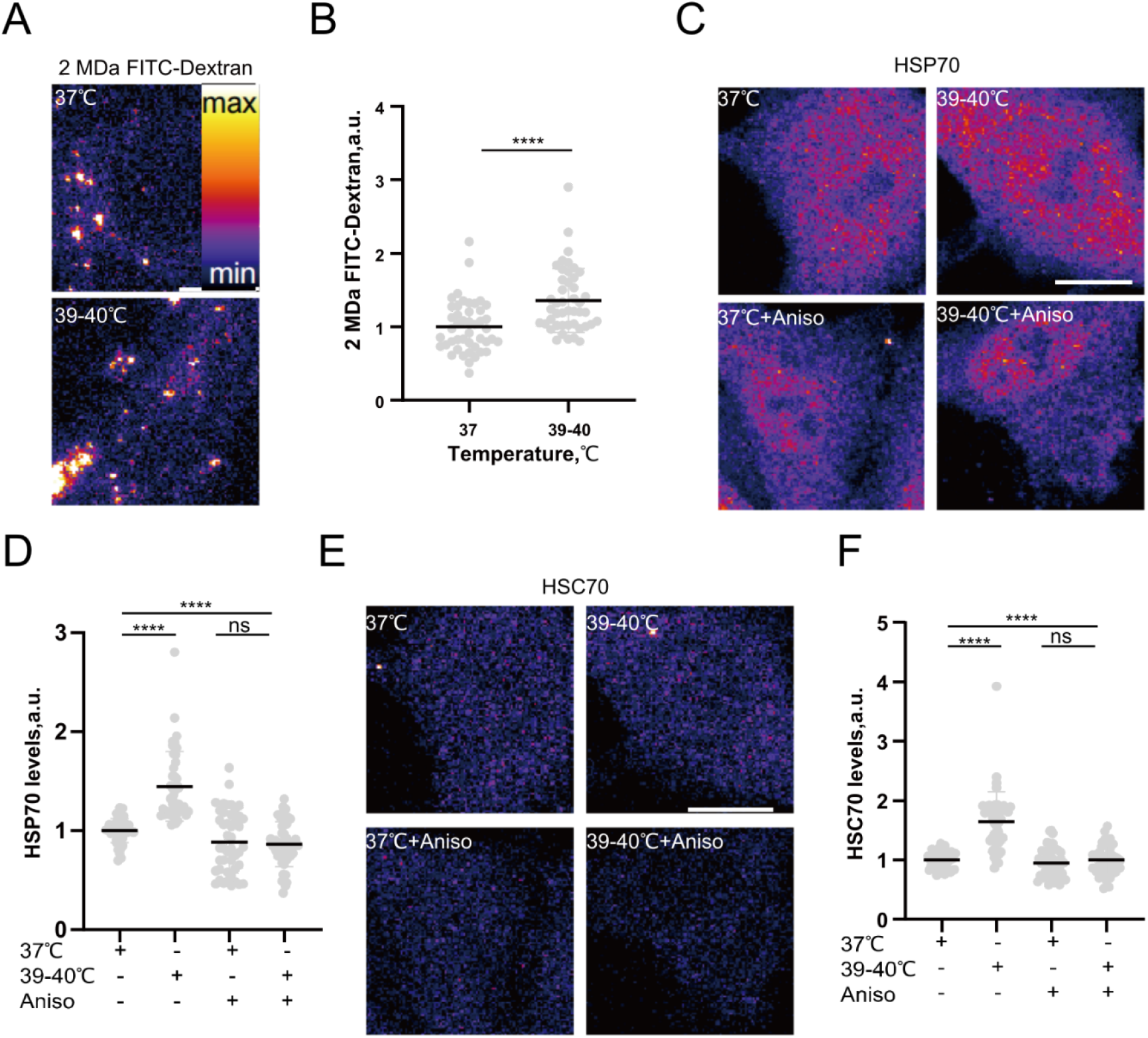
Hyperthermia triggers heat shock response. (A) Representative images of 2 MDa FITC-dextran internalisation in PC12 cells subjected to hyperthermia for 3 hours. (B) Quantification of 2 MDa FITC-dextran endocytosis.****P < 0.0001, Mann–Whitney test. (C) Representative images of HSP70 in PC12 cells treated with hyperthermia for 3 hours and anisomycin. (D) Quantification of HSP70 levels in PC12 cells treated with hyperthermia for 3 hours and anisomycin. ****P < 0.0001, ns-not significant, Kruskal–Wallis test with Dunn’s multiple comparisons test. (E) Representative images of HSC70 levels in PC12 cells treated with hyperthermia for 3 hours and anisomycin. (F) Quantification of HSC70 levels in PC12 cells treated with hyperthermia for 3 hours and anisomycin.****P < 0.0001, ns-not significant, Kruskal–Wallis test with Dunn’s multiple comparisons test. N ≥ 3 independent experiments, n = 45 cells. Scale bar, 10 μm.

**Figure S3.**
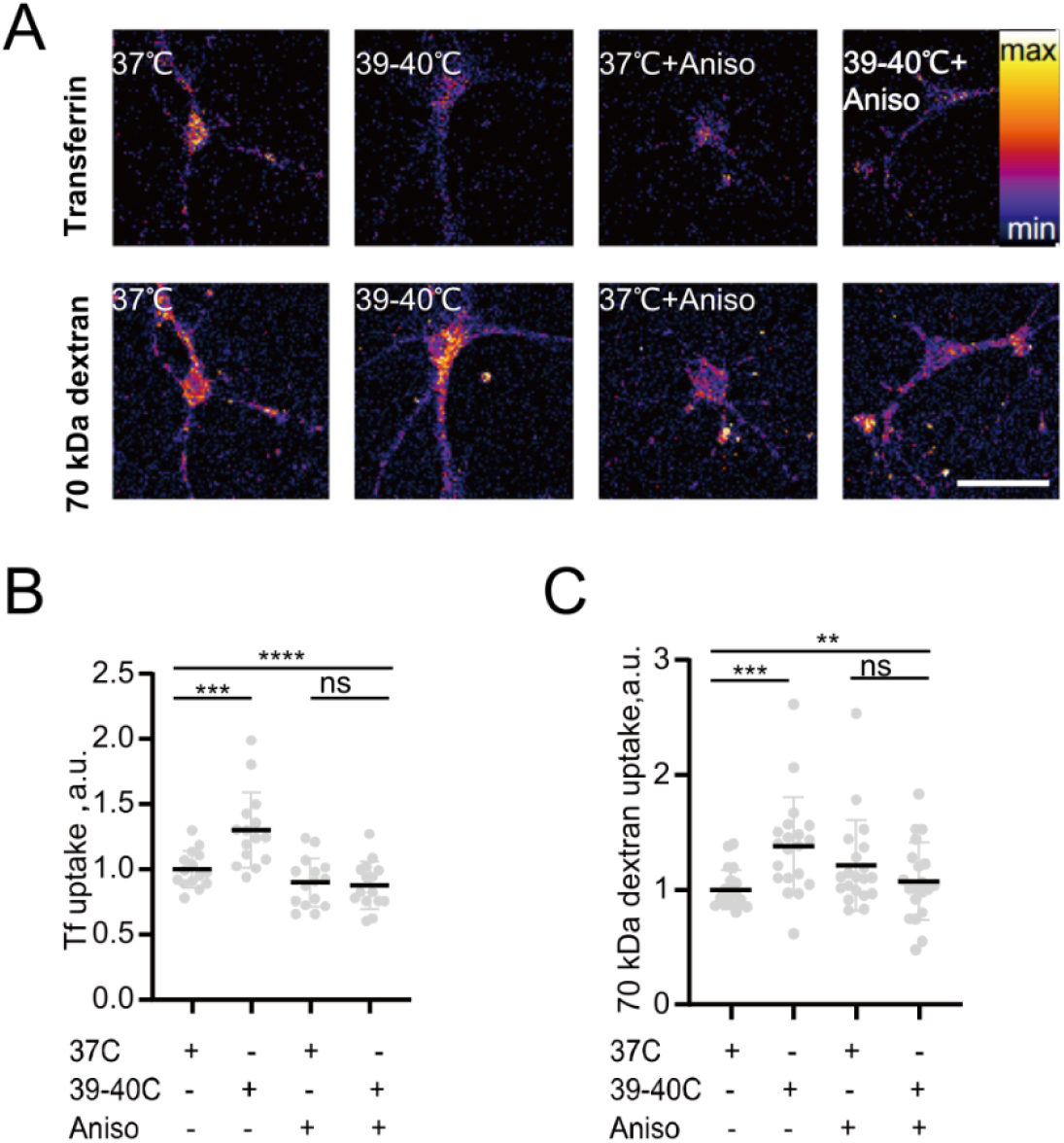
Hyperthermia triggers membrane trafficking in primary neurons. (A) Representative images of FITC-Tf and 70 kDa TRITC-dextran internalisation in primary neurons treated with hyperthermia for 3 hours and anisomycin. (B) Quantification of FITC-Tf internalisation in primary neurons treated with hyperthermia for 3 hours with 5 anisomycin. ****P < 0.0001,***P < 0.001, ns-not significant, Kruskal–Wallis test with Dunn’s multiple comparisons test. (C) Quantification of 70 kDa TRITC-dextran internalisation in primary neurons treated with hyperthermia for 3 hours with anisomycin.***P < 0.001,**P < 0.01, ns-not significant, Kruskal–Wallis test with Dunn’s multiple comparisons test. N ≥ 3 independent experiments, n ≥ 15 cells. Scale bar, 50 μm.

**Figure S4.**
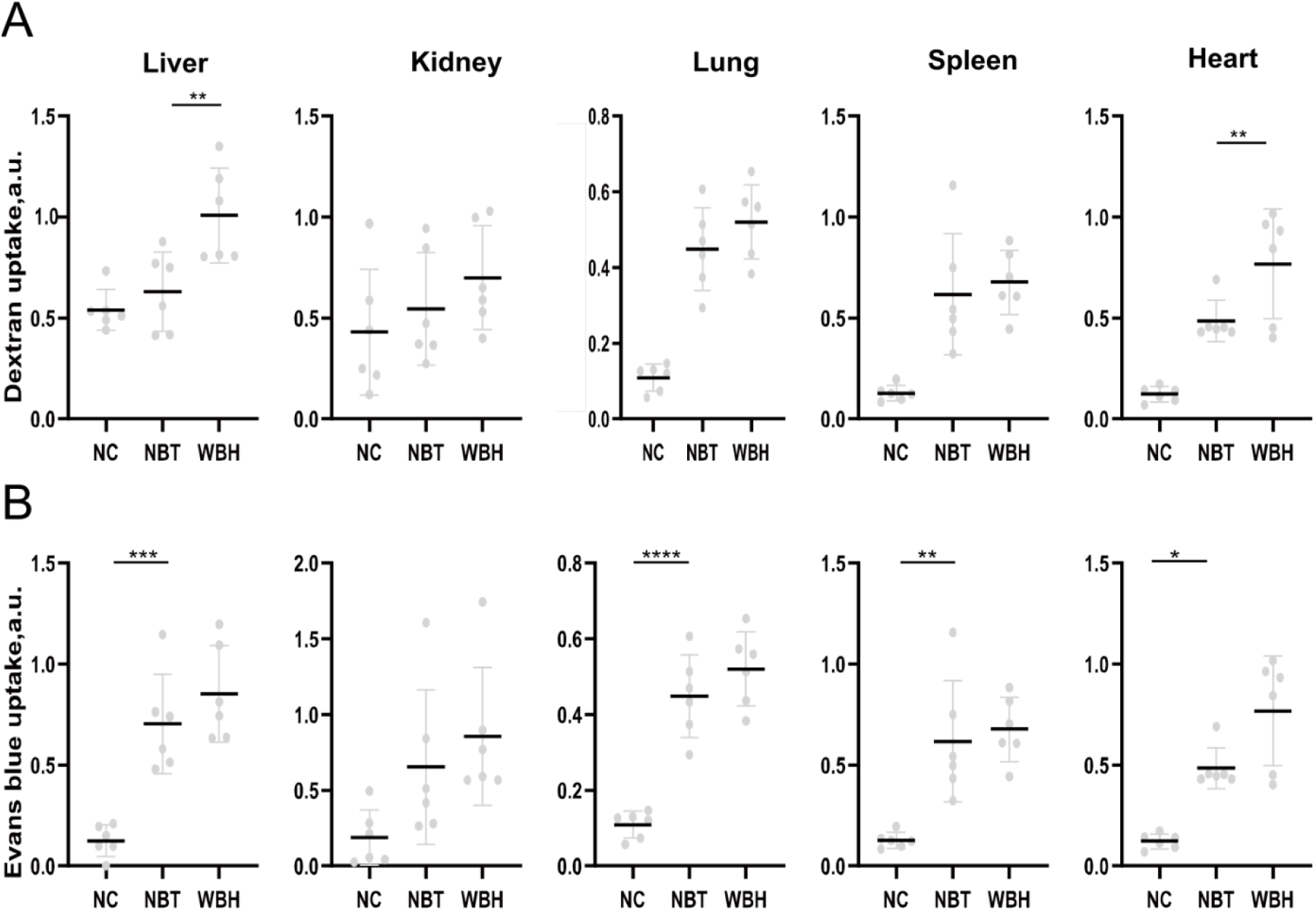
Effects of whole body mild hyperthermia on fluid phase endocytosis across the body. (A) Quantification of FITC signal in organs of mice injected intraperitoneally 24 hours prior to 6 hours of WBH. (B) Quantification of Evans blue in organs of mice injected intraperitoneally 24h prior to 6 hours of WBH. ****P<0.0001, **P<0.01, Kruskal–Wallis test with Dunn’s multiple comparisons test. 6 animals/condition.

**Table S1.** Antibodies used in this study.

| Antigen | Species | Conjugation | Manufacturer | Cat. No. | Dilution |
| --- | --- | --- | --- | --- | --- |
| EEA1 | rabbit |  | Abcam | ab2900 | 1/300 |
| LAMP1 | mouse |  | Invitrogen | MA1-164 | 1/300 |
| Rabbit IgG | goat | Alexa Fluor 488 | Bioss | Bs-0295<br>G-AF488 | 1/500 |
| Mouse IgG | goat | Alexa Fluor 594 | SolarBio | K1031G-AF<br>594 | 1/500 |
| HSP70 | rabbit |  | Boster | BA0928 | 1/400 |
| HSC70 | mouse |  | Boster | M01024-1 | 1/400 |
| GABAAR | rabbit | | Synaptic Systems | 224203 | 20 μg/kg |

